# RNA-seq sample preparation kits strongly affect transcriptome profiles of a gas-fermenting bacterium

**DOI:** 10.1101/2022.04.28.489910

**Authors:** Lorena Azevedo de Lima, Kristina Reinmets, Lars Keld Nielsen, Esteban Marcellin, Audrey Harris, Michael Köpke, Kaspar Valgepea

**Author notes:** Address correspondence to Kaspar Valgepea. These authors contributed equally to this work (Lorena Azevedo de Lima, Kristina Reinmets). Author order was determined alphabetically.

## Abstract

Transcriptome analysis via RNA sequencing (RNA-seq) has become a standard technique employed across various biological fields of study. This rapid adoption of the RNA-seq approach has been mediated, in part, by the development of different commercial RNA-seq library preparation kits compatible with standard next-generation sequencing (NGS) platforms. Generally, the essential steps of library preparation such as ribosomal RNA (rRNA) depletion and first-strand cDNA synthesis are tailored to a specific group of organisms (e.g. eukaryotes vs. prokaryotes) or genomic GC content. Therefore, the selection of appropriate commercial products is of crucial importance to capture the transcriptome of interest as closely to the native state as possible without introduction of technical bias. However, researchers rarely have the resources and time to test various commercial RNA-seq kits for their samples. This work reports a side-by-side comparison of RNA-seq data from *Clostridium autoethanogenum* obtained using three commercial rRNA removal and strand-specific library construction products by NuGEN Technologies, Qiagen, and Zymo Research and assesses their performance relative to published data. While all three vendors advertise their products as suitable for prokaryotes, we found significant differences in their performance regarding rRNA removal, strand-specificity, and, most importantly, transcript abundance distribution profiles. Notably, RNA-seq data obtained with Qiagen products were most similar to published data and delivered the best results in terms of library strandedness and transcript abundance distribution range. Our results highlight the importance of finding appropriate organism-specific workflows and library preparation products for RNA-seq studies.

**Importance:** RNA-seq is a powerful technique for transcriptome profiling while involving elaborate sample processing before library sequencing. Our work is important as we show that RNA-seq library preparation kits can strongly affect the outcome of the RNA-seq experiment. Although library preparation benefits from the availability of various commercial kits, choosing appropriate products for the specific samples can be challenging for new users or for users working with unconventional organisms. Evaluating the performance of different commercial products requires significant financial and time investment infeasible to most researchers. Therefore, users are often guided in their choice of kits by published data involving similar input samples. We conclude that important consideration should be given to selecting of sample processing workflows for any given organism.

## Introduction

Gene transcription is a fundamental process mediating vast number of intracellular and environmental responses in every cell. Therefore, understanding transcriptional states of any organism of choice can shed light on basic biological processes as well as ways to direct and control cellular behavior. Insights into cellular transcriptional profiles or transcriptomes (i.e. complete set of transcripts in a cell with their quantities) have vastly expanded in the last decade due to rapid development and high accessibility of next generation sequencing (NGS) platforms (1). Meanwhile, constant improvement of commercial library construction products has greatly contributed to the rapid adaptation and evolution of RNA sequencing applications: for example, RNA-seq (2), nascent RNA sequencing (3), Ribo-seq (4), and differential RNA-seq (5). RNA-seq is the most common application as it allows both mapping and quantification of transcriptomes.

While RNA-seq has become widely used across all fields of biological sciences, obtaining high-quality data of the transcriptome under investigation nevertheless requires careful planning, extensive sample processing, and considerable resources. The availability of commercial RNA-seq library preparation kits tailored to a variety of organisms, experimental approaches, and sequencing platforms has made RNA-seq accessible even to non-expert users. When planning to do an RNA-seq experiment for the first time, researchers often consult existing literature to see which sample preparation protocols and products have been previously used with their organism of interest. However, working with unconventional microorganisms that have not yet been extensively studied via RNA-seq can make it difficult to decide which commercial kits might be most suitable.

We have previously achieved high-quality transcriptome profiling using RNA-seq for the gas-fermenting bacterium *Clostridium autoethanogenum* (6–8), an unconventional microbe that is also used as a cell factory in commercial-scale gas fermentation for the production of low carbon fuels and chemicals from waste feedstocks (9). In addition to preparation of cDNA libraries before sequencing, removal of ribosomal RNA (rRNA) from the extracted total RNA is needed to ensure efficient transcriptome-wide messenger RNA (mRNA) detection and quantification as >80−90% of total cellular RNA is rRNA (10, 11). In our previous studies (6–8), we used Illumina kits for rRNA removal and library preparation but when we set out to start a large-scale RNA-seq survey of the same organism in late 2019, the Illumina Ribo-Zero rRNA removal kit was discontinued, and we had to look for alternatives. However, selecting an efficient rRNA removal method for bacterial samples is non-trivial as enrichment of non-rRNA transcripts based on polyadenylated RNA (polyA) selection used in most commercial kits (developed for eukaryotes) is not applicable for bacterial RNA due to the lack of polyA tails. One also has to ensure the compatibility of the rRNA removal and cDNA library preparation methods.

To make an informed decision on the following best commercial products for RNA-seq library preparation for *C. autoethanogenum*, we aimed to test kits from three vendors that are advertised to ensure efficient rRNA removal and to be compatible with a variety of bacterial species and Illumina sequencing platforms. This work reports a side-by-side comparison of RNA-seq data obtained from the same *C. autoethanogenum* input samples using rRNA removal and strand-specific library construction kits from NuGEN Technologies, Qiagen, and Zymo Research, and assesses their performance relative to published data. Transcriptome profiles revealed significant differences between the kits regarding rRNA removal efficiency, sequencing reads strand-specificity, and, strikingly, in transcript abundance distribution profiles. Our work shows that Qiagen kits yield the most reliable data out of the three we tested and highlights the importance of appropriate sample preparation for RNA-seq analysis in bacteria.

## Results and Discussion

### Experimental design

We evaluated the performance of three commercial rRNA removal and strand-specific library construction kits by NuGEN, Qiagen, and Zymo (see Materials and Methods for details) for RNA-seq analysis of *C. autoethanogenum* autotrophic cultures (Fig. 1A). To assess the ability of the selected commercial kits to capture the transcriptomic profile of *C. autoethanogenum* under varying culture conditions, we used four samples, each obtained from one of the four bioreactor continuous culture experiments grown on two different feed gas mixes (CO or CO+CO_2_+H_2_ [syngas]) and dilution rates (i.e. specific growth rates; 1 or 2 day^-1^). Both feed gas composition (12) and specific growth rate (13) of the culture have profound effects on the culture phenotype (e.g. gas uptake, product distribution, metabolic fluxes). We extracted and prepared total RNA from the four samples using a previously established workflow optimised for *C. autoethanogenum* (6). Next, total RNA for each sample was split between the NuGEN, Qiagen, and Zymo kits for rRNA removal and strand-specific RNA-seq library construction according to vendors’ instructions. Finally, the 12 samples (four cultures times three kits) were examined by paired-end 75 bp sequencing on an Illumina MiSeq platform, followed by RNA-seq data analysis using established pipelines (6, 13).

**FIG 1.**
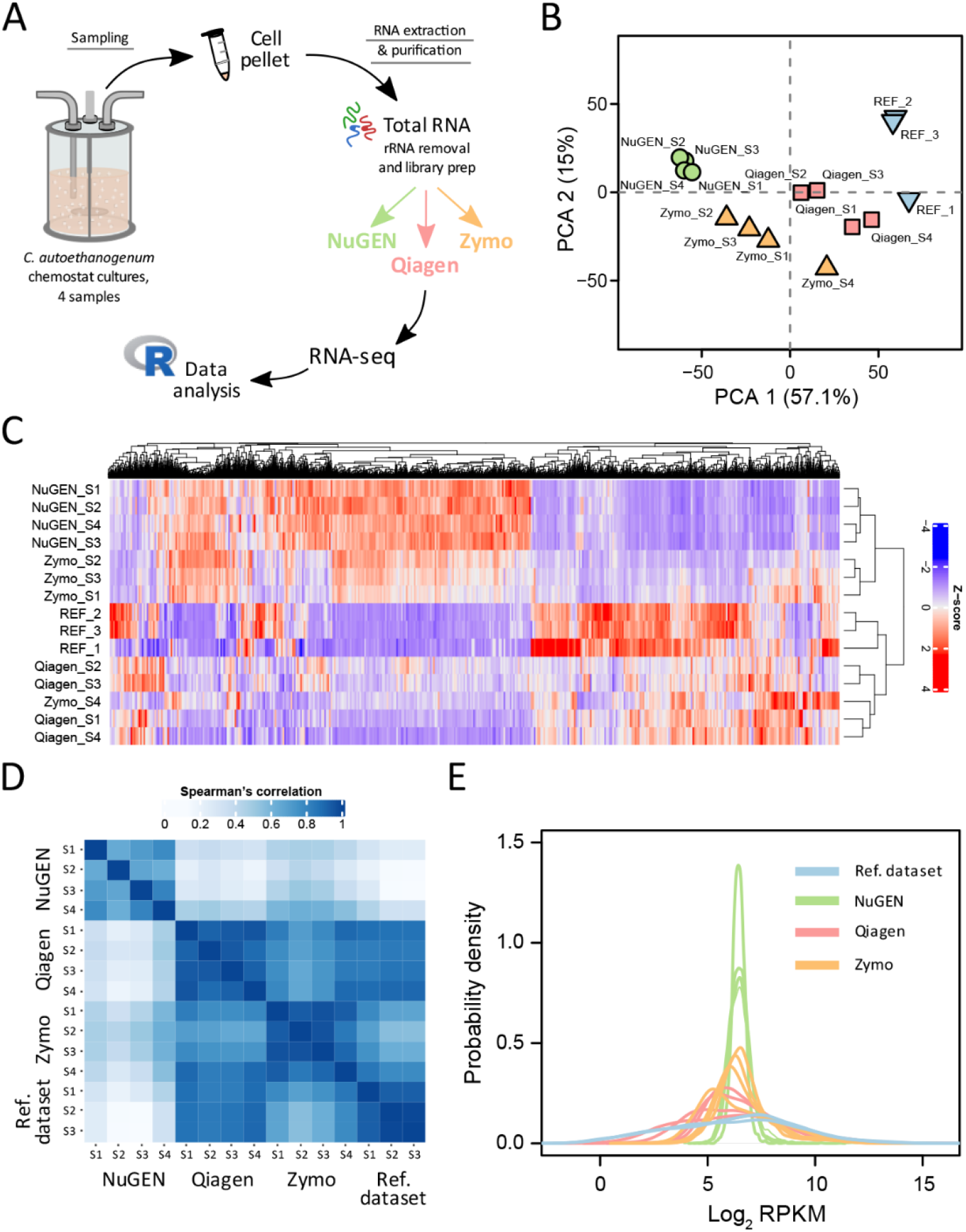
RNA-seq results are strongly affected by rRNA removal and library construction kits. (A) Experimental design of this work. (B) Principal component analysis (PCA) of transcript abundances. (C) Hierarchical clustering of individual transcript abundances. (D) Spearman’s correlation analysis of transcript abundances. (E) Probability density plots of transcript abundances. The reference dataset refers to high BC samples in GEO accession number GSE90792. REF, reference dataset. rRNA transcript abundances were removed prior to data analysis to avoid bias from variable efficiency of rRNA removal between kits.

### General statistics of RNA-seq data

An average of 4.5 million raw reads per sample were obtained from the sequencing runs that were mapped to the reference genome of *C. autoethanogenum* NC_022592.1 (14) after trimming with an overall high success rate (Table 1). Namely, 98, 93, and 99% of reads were mapped on average for NuGEN, Qiagen, and Zymo, respectively, which resulted in a minimum of 50-fold coverage of the *C. autoethanogenum* genome across samples (Table 1). We detected very low read duplication levels (<0.5%) suggesting a low chance of technical bias introduced during sample preparation. Surprisingly, a significant difference in the percentage of mapped reads that were assigned to genomic features (i.e. FeatureCounts) was observed between NuGEN and the other two kits: an average of only 55% for NuGEN with 84 and 79% for Qiagen and Zymo, respectively (Table 1; FeatureCounts/Mapped). Notably, this can be explained by the difference between NuGEN and the other two kits in the percentage of reads mapping to the expected strand, as Qiagen and Zymo showed high correct strandedness at ∼91 (sense) and ∼84% (antisense), respectively, compared to NuGEN’s very poor strand-specificity at ∼58% (sense) (Table 1). Significant false strandedness for NuGEN could arise from either a substantial flaw in the respective workflow or, according to NuGEN, from faulty reagents in their kits (personal communication).

**TABLE 1.** General statistics of RNA-seq results of the three tested kits for rRNA removal and library construction

| Kits | Sample name | Number of raw reads | Reads mapped (Mapped) | Coverage (fold) | FeatureCounts /Mapped | Strandedness |  | rRNA RPKM/ Total RPKM |
| --- | --- | --- | --- | --- | --- | --- | --- | --- |
|  |  |  |  |  |  | Sense | Antisense |  |
| NuGEN | NuGEN_S1 | 4,268,524 | 98% | 72 | 59% | 64% | 36% | 7% |
|  | NuGEN_S2 | 6,004,018 | 99% | 103 | 47% | 50% | 50% | 2% |
|  | NuGEN_S3 | 3,615,864 | 99% | 62 | 52% | 56% | 44% | 4% |
|  | NuGEN_S4 | 4,248,288 | 98% | 72 | 63% | 61% | 39% | 15% |
| Qiagen | Qiagen_S1 | 4,911,456 | 98% | 81 | 87% | 95% | 5% | 15% |
|  | Qiagen_S2 | 3,559,522 | 95% | 57 | 79% | 84% | 16% | 9% |
|  | Qiagen_S3 | 3,289,702 | 93% | 50 | 82% | 88% | 12% | 6% |
|  | Qiagen_S4 | 4,954,218 | 88% | 74 | 88% | 96% | 4% | 17% |
| Zymo | Zymo_S1 | 4,355,956 | 99% | 71 | 82% | 13% | 87% | 0.8% |
|  | Zymo_S2 | 4,840,762 | 99% | 79 | 70% | 26% | 74% | 0.6% |
|  | Zymo_S3 | 5,691,744 | 99% | 93 | 78% | 18% | 82% | 0.6% |
|  | Zymo_S4 | 4,651,118 | 99% | 76 | 87% | 7% | 93% | 0.8% |

rRNA, ribosomal RNA; RPKM, reads per kilobase of transcript per million mapped reads.

### Variable efficiency of rRNA removal

We next quantified rRNA removal efficiencies from the RNA-seq data using the percentage of rRNA transcript abundances from total transcript abundances, expressed as reads per kilobase of transcript per million mapped reads (RPKM) (Table S1). Again, stark differences between the kits were observed, confirming that rRNA depletion from bacterial samples is non-trivial (Table 1, rRNA RPKM/Total RPKM). Zymo demonstrated superior efficiency for rRNA removal during library preparation with an abundance of <1% of rRNA transcripts. NuGEN’s higher variability in rRNA removal efficiency across the samples (2−15%; average 7%) suggests their approach of non-rRNA enrichment or AnyDeplete™ technology may be sensitive to sample-specific factors. Removal of rRNA for Qiagen was slightly less efficient (∼11%) than for NuGEN but still acceptable to ensure high coverage of transcriptome-wide mRNA detection and quantification (Table 1).

### Kit-specific grouping of transcriptome profiles

Upon observing the differences in general RNA-seq metrics outlined in Table 1, we were curious if different kits could also lead to variable transcriptome profiles. Indeed, principal component analysis (PCA) of transcript abundances revealed clear sample grouping by the kit and not by the origin of the input RNA (Fig. 1B). To assess which of the three tested kits produced the most reliable transcriptome profiles, we also included published data in the PCA that we previously obtained using the same workflows but with Illumina kits for similar *C. autoethanogenum* culture conditions (6), termed here as the reference dataset (high biomass concentration samples in GEO accession number GSE90792). Notably, Qiagen data was grouped the closest to this reference dataset with NuGEN transcriptome profiles separating most distinctively (Fig. 1B). These observations were confirmed by hierarchical clustering of individual transcript abundances showing grouping of samples based on the kits and not based on the origin of the input RNA (Fig. 1C). NuGEN and Zymo had a distinctively different clustering pattern compared to Qiagen and the reference dataset.

Clustering results agreed with Spearman’s correlation analysis of transcript abundances between samples, which showed Qiagen data being most similar to the reference dataset (ρ ∼ 0.86) (Fig. 1D). Within the three kits tested here, Qiagen and Zymo data showed higher similarity (ρ ∼ 0.77 across the same samples) compared to the lower correlations between NuGEN and Qiagen (ρ ∼ 0.33) and NuGEN and Zymo (ρ ∼ 0.47) data.

### Differences in transcript abundance distribution profiles

The quality of the kits can also be assessed by their sensitivity to detect transcripts across a range of abundances (i.e. transcriptome coverage or depth). Transcript levels in bacteria generally span over 4 orders of magnitude (15–17), including in *C. autoethanogenum* (6) and other gas-fermenting bacteria (18, 19). Again, Qiagen data resembled the reference dataset the most by both the transcript abundance distribution profiles and good sensitivity with transcript levels spanning over 4 orders of magnitude (from ∼2 to ∼39,000 RPKM; ∼1 to 15 log_2_ RPKM) (Fig. 1E). Strikingly, NuGEN kit showed very narrow transcript abundance distributions covering only ∼2 orders of magnitude (from ∼22 to ∼1,800 RPKM; ∼4 to ∼11 log_2_ RPKM), while Zymo data were positioned between Qiagen and NuGEN. According to Zymo (personal communication), such condensed distribution profiles could be caused by higher sensitivity of the workflow towards the presence of genomic DNA in the input sample that can artificially inflate mRNA reads with a more prominent effect on low-abundance transcripts, thereby pushing the left tail of the distribution to the right. This would also be consistent with the poorer strandedness of Zymo and NuGEN data (Table 1) arising from genomic DNA-originating reads. Our sample preparation workflow previously optimised for *C. autoethanogenum* (6) efficiently removed DNA from total RNA samples down to ∼13 ± 2 ng/μL (average ± standard deviation), making up ∼4% of the RNA concentration. Thus, additional steps to deplete DNA to extremely low levels are potentially required for Zymo and NuGEN workflows. NuGEN data could be additionally explained by biased synthesis and amplification of cDNA using selective primers compared to the general use of random primers in RNA-seq workflows.

Our work is important as researchers rarely have the resources and time to test various commercial RNA-seq kits, advertised as suitable for multiple organisms with different genomic GC content, for their samples. The ability to capture the spectrum of transcript abundances as closely to the true cellular state as possible is crucial to accurately address research questions investigated via RNA-seq. Our work shows that rRNA removal and library construction kits can strongly affect RNA-seq outcomes. This is highly relevant for anyone establishing an RNA-seq pipeline for an organism or for researchers puzzled by unexpected RNA-seq results. We conclude that, at least for *C. autoethanogenum* RNA-seq studies, Illumina and Qiagen kits are most suitable by providing high sensitivity across a wide range of transcript levels, superior strand-specificity, and sufficient rRNA removal, ensuring high coverage of transcriptome-wide mRNA detection and quantification.

## Materials and Methods

### Bacterial strain and cultivation conditions

A derivate of *Clostridium autoethanogenum* DSM 10061 strain – DSM 23693 – deposited in the German Collection of Microorganisms and Cell Cultures (DSMZ) was used in all experiments and stored as a glycerol stock at -80°C. Full details of the cultivation conditions are reported in previous work (13). Shortly, cells were grown autotrophically in bioreactor chemostat continuous cultures under strictly anaerobic conditions at 37°C and pH 5 in chemically defined medium (without yeast extract) either on CO (60% CO and 40% Ar; AS Eesti AGA) or syngas (50% CO, 20% H_2_, 20% CO_2_, and 10% Ar; AS Eesti AGA). Namely, four independent experiments were conducted with cultures grown at dilution rates (D) ∼1.0 and ∼2.0 day^-1^ on both feed gas mixes. Cultures were sampled for RNA extraction and subsequent transcriptome analysis using RNA-seq after optical density (OD), gas uptake, and production rates had been stable for at least one working volume.

### Preparation of total RNA extracts

Full details of culture sampling, RNA extraction, and purification are reported in previous work (13). Briefly, culture samples were pelleted by centrifugation and treated with RNAlater (76106; Qiagen) before disrupting cells with glass beads using the Precellys® 24 instrument and extracting total RNA using the RNeasy Mini Kit (74104; Qiagen). Next, RNA extracts were depleted of DNA using off-column TURBO™ DNase (AM2239; Invitrogen) followed by purification using the RNA Clean & Concentrator™ kit (R1018; Zymo). We used the NanoDrop™ 1000 instrument (Thermo Fisher Scientific) for verifying efficiency of RNA purification. The high quality and integrity of the total RNA extracts was confirmed by RNA integrity numbers (RIN) above 8.2 using the TapeStation 2200 equipment (Agilent Technologies). Total RNA and residual DNA concentrations were determined using the Qubit 2.0 instrument (Invitrogen).

### Removal of rRNA and RNA-seq library construction

Total RNA extracts for each sample were split between the NuGEN, Qiagen, and Zymo kits for rRNA removal and strand-specific RNA-seq library construction according to vendor instructions. In this work, samples referred to as “NuGEN” were processed with Universal Prokaryotic RNA-Seq, Prokaryotic AnyDeplete™ (0363; NuGEN); “Qiagen” with QIAseq® FastSelect™ –5S/16S/23S Kit (335925; Qiagen) (for rRNA removal) and QIAseq® Stranded RNA Lib Kit (180743; Qiagen) (for library construction); and “Zymo” with Zymo-Seq RiboFree™ Total RNA Library Kit (R3000; Zymo Research).

### RNA sequencing and data analysis

RNA sequencing of the 12 mRNA libraries (four cultures times three kits) was performed on a MiSeq instrument (Illumina) using the MiSeq v3 150 cycles sequencing kit (MS-102-3001; Illumina) with paired-end 2 × 75 bp reads. Raw RNA-seq data of the reference dataset (high biomass concentration samples in GEO accession number GSE90792) (6) was analysed together with the data generated in this work to ensure comparability. Full details of RNA-seq data analysis, including R-scripts, are reported in previous work (13). Shortly, quality of sequencing reads was verified using MultiQC (20) and presence of read duplicates was examined using PicardTools (21). High-quality reads were then mapped to the NCBI reference genome of *C. autoethanogenum* NC_022592.1 (14) and genomic features were assigned using Rsubread (22). Strandedness of reads for the strand-specific data of NuGEN, Qiagen, and Zymo was calculated using RSeQC v3.0.1 (23). Lastly, raw library sizes were normalised and transcript abundances were estimated as reads per kilobase of transcript per million reads mapped (RPKM) using edgeR (24) (see Table S1 for RPKM data). rRNA transcript abundances were removed prior to data analysis on Fig. 1 to avoid bias from variable efficiency of rRNA removal between kits. Hierarchical clustering of individual transcript abundances on Fig. 1C was performed using the ComplexHeatmap package in R (version 2.10.0) (25). RNA-seq data has been deposited in the NCBI Gene Expression Omnibus repository under accession number GSE200959.

## Acknowledgements

We thank Viktorija Kukuškina for valuable discussions.

This work was funded by the European Union’s Horizon 2020 research and innovation programme under grant agreement N810755 and the Estonian Research Council’s grant agreement PSG289. Australian Government funding through its investment agency, the Australian Research Council, towards the ARC Centre of Excellence in Synthetic Biology (CE200100029) is gratefully acknowledged. We thank the following investors in LanzaTech’s technology: BASF, CICC Growth Capital Fund I, CITIC Capital, Indian Oil Company, K1W1, Khosla Ventures, the Malaysian Life Sciences, Capital Fund, L. P., Mitsui, the New Zealand Superannuation Fund, Novo Holdings A/S, Petronas Technology Ventures, Primetals, Qiming Venture Partners, Softbank China, and Suncor.

LanzaTech has interest in commercial gas fermentation with *C. autoethanogenum*. AH and MK are employees of LanzaTech.

